# Astigmatism-based focus stabilisation with universal objective lens compatibility, extended operating range and nanometre precision

**DOI:** 10.1101/2024.01.15.575442

**Authors:** Amir Rahmani, Tabitha Cox, Akhila Thamaravelil Abhimanue Achary, Aleks Ponjavic

## Abstract

Focus stabilisation is vital for long-term fluorescence imaging, particularly in the case of high-resolution imaging techniques. Current stabilisation solutions either rely on fiducial markers that can be perturbative, or on beam reflection monitoring that is limited to high-numerical aperture objective lenses, making multimodal and large-scale imaging challenging. We introduce a beam-based method that relies on astigmatism, which offers advantages in terms of precision and the range over which focus stabilisation is effective. This approach is shown to be compatible with a wide range of objective lenses (10x-100x), typically achieving <10 nm precision with >10 μm operating range. Notably, our technique is largely unaffected by pointing stability errors, which in combination with implementation through a standalone Raspberry Pi architecture, offers a versatile focus stabilisation unit that can be added onto most existing microscope setups.

## Introduction

Optical microscopy typically relies on imaging of samples positioned at the focal plane of the microscope objective lens. Real-time maintenance of the in-focus imaging plane is crucial due to potential axial drift caused by external factors. These include mechanical instabilities and thermal influences ^1^ that are particularly problematic for long-term imaging, where instabilities can degrade the imaging quality. ^2^ With the development of super-resolution microscopy techniques that achieve nanoscale imaging, ^3–7^ it is vital to implement active focus stabilisation, especially in the case of high-throughput automated microscopy. ^8–11^

Numerous focus stabilisation approaches have been developed for applications in optical microscopy. ^12,13^ Building on these, several commercial systems have been designed that can maintain focus during imaging. ^14–17^ However, their limited compatibility with custom-built microscopes, and their significant cost, has hampered their adoption. This has resulted in a variety of custom approaches that can be classified as: (1) marker-based ^18–20^ or (2) beam-based methods (Fig. 1a). ^21–26^ In the marker-based approach, the fluorescence or scattering from bright particles is collected through a dedicated channel distinct from the imaging channel. ^27,28^ By monitoring these fiducial markers, their drift can be corrected. Recently, this approach has enabled fast active 3D-focus stabilisation by integrating GPU processing. ^29^ However, the introduction of fiducial markers is: (1) not always available depending on the samples; (2) has the potential to influence the sample of interest and (3) adds an additional experimental step. By using a beam-based approach instead, these issues can be avoided.

**Fig. 1.**
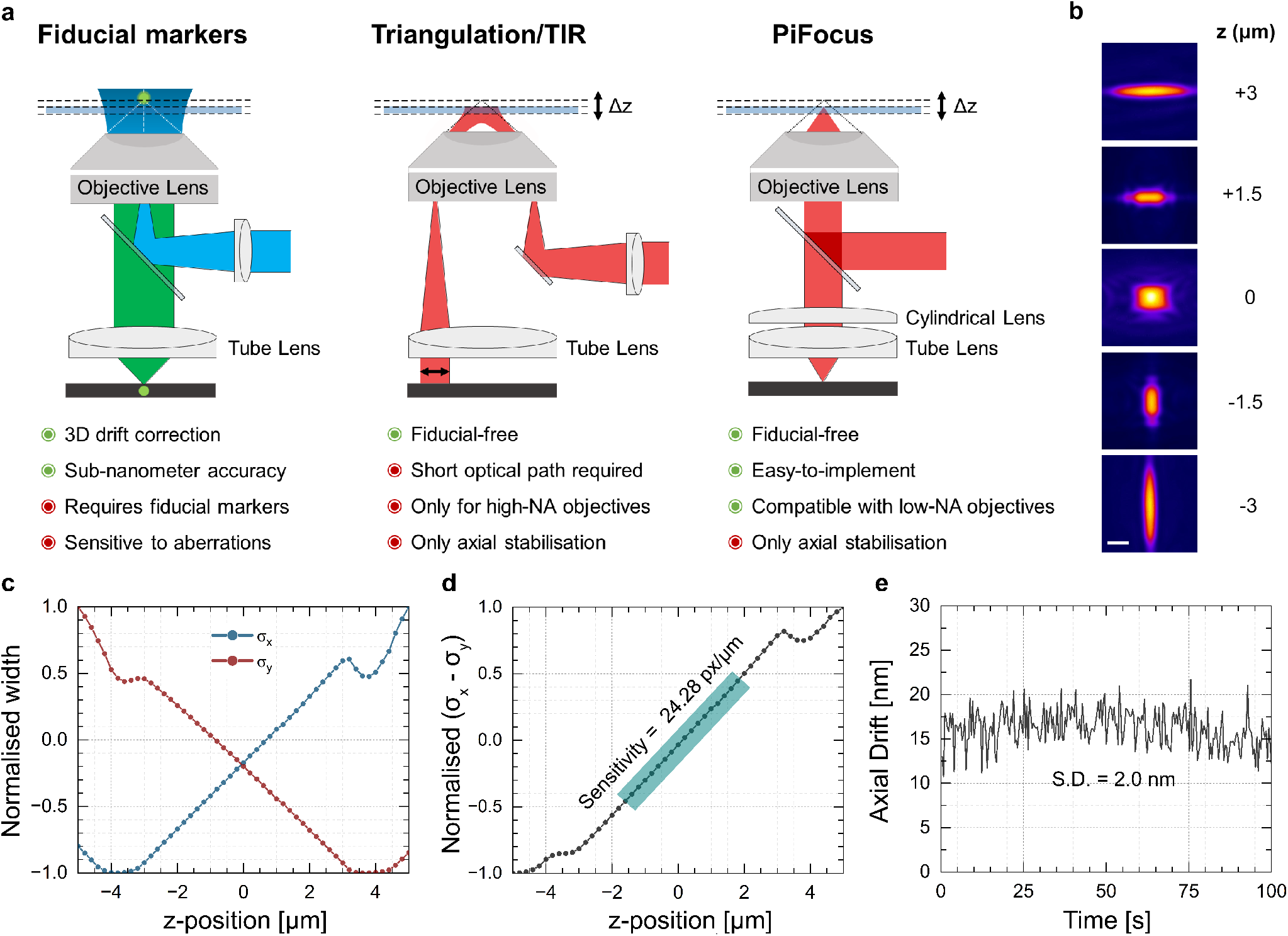
Axial drift estimation using astigmatism. (a) Schematic and characteristics of common focus stabilisation techniques including PiFocus presented in this work. (b) Selected slices from a z-scan of the astigmatic PSF (scale bar = 40 pixels). (c) Variation of beam widths of *σ*_*x*_ (blue line) and *σ*_*y*_ (red line) as a function of z-position. (d) Calibration curve of focal drift estimation using *σ*_*x*_ −*σ*_*y*_. (e) Time-lapse acquisition at z = 0 μm for estimating axial precision.

With the beam-based approach, it is common to monitor the reflection of an infrared laser from the coverslip-sample interface. Among beam-based methods, the triangular geometry or total internal reflection (TIR) has been widely adopted, and methods that incorporate open hardware and comprehensive documentation have been developed. ^30^ Nevertheless, there are limitations of current approaches that can be summarised by: (1) relatively low sampling rates (pgFocus: 30 Hz, ^31^ openFrame: 4 Hz ^32^); (2): only being compatible with high numerical aperture (NA); (3) electronic components requiring soldering and custom circuit boards and (4): the need to put a focus stabilisation module close to the objective lens as the performance is limited by laser pointing stability. ^33^ Alternative beam-based methods have recently been introduced, e.g. parallel beam splitting by a double-hole mask. ^34^ However, these non-TIR approaches still suffer from low precision in axial drift estimation and stabilisation. ^35–39^ An ideal focus stabilisation system that would improve on existing iterations would be: (1) standalone, (2) open-source, (3) low cost, (4) fiducial-free and (5) would offer large range and nanoscale precision with active stabilisation.

A promising solution involves applying astigmatism for focus stabilisation ^40,41^ using a reflected beam from the coverslip-sample interface. Its widespread application may have been impeded by the interference patterns generated from reflective surfaces due to the use of coherent laser sources. ^42^ However, this issue has recently been resolved in the scattering microscopy field ^43,44^ and in a recent focus stabilisation method. ^32^ Here we present PiFocus, a versatile focus stabilisation method implemented on Raspberry Pi, based on monitoring the astigmatic reflection from the coverslip-sample interface. We demonstrate that this method enables <10 nm axial drift control with extended operating range (>10 μm). Excitingly, it becomes possible to tune the operating range and precision such that ∼20 nm precision can be achieved even with low-magnification (10x), low-NA objective lenses. Ultimately, PiFocus is implemented as a low-cost (<£2,000) standalone system that can be added on to most microscopes, with advantages such as ease of application, wide objective lens compatibility and versatility.

## Results

### Astigmatic-based drift estimation with nanometre precision

Our proposed focus stabilisation approach relies on determining the axial position of the sample by inducing astigmatism to the back-reflected laser beam from the coverslip-sample interface. PiFocus was initially implemented using a 100x 1.35-NA silicone-immersion objective lens and astigmatism was induced with a cylindrical lens placed in front of the camera (Fig. 1a). The point-spread-function (PSF) is symmetric when the sample is in focus and becomes elliptical when defocused (Fig. 1b). By measuring the beam widths in x and y (Fig. 1c), here referred to as *σ*_*x*_ and *σ*_*y*_, the axial sample position and the direction of its displacement can be found based on a calibration scan (Fig. 1d, Video S1). In each axial step the difference between *σ*_*x*_ and *σ*_*y*_ is determined. We used a multimode infrared laser (808 nm) to ensure minimal overlap with common fluorescence channels. The multimode laser provided a broadband beam (coherence length of 32 μm and bandwidth 9 nm), which avoids the interference pattern that would otherwise be created due to secondary reflections in the beam path (Fig. S1, Video S2). ^32,43^ This laser was delivered through a single-mode fibre to provide a Gaussian beam that could be shaped for axial drift estimation. Some experiments (TIR, 40x and 10x) used a 638 nm multimode laser instead for simpler alignment, but the results were identical (Fig. S2).

To determine the accuracy of this approach, a time-lapse (100 Hz) acquisition was collected at a specific axial position (z = 0 μm). The duration of the time-lapse was typically a few seconds to minimise drift. The uncertainty of *σ*_*x*_ − *σ*_*y*_ of the time-lapse was then converted to nanometres using the calibration (Fig. 1d), to estimate the precision of the approach (Fig. 1e), which was found to be *σ*_*z*_ = 2.0 nm for these conditions. Note that this precision is valid at z = 0 and we observe an approximately two-fold increase in the precision value towards the limits of the defined operating range, as is common for astigmatic localisation microscopy. ^45^

### Tuning the focus stabilisation operating range

We next investigated the ability of PiFocus to improve the axial range over which the focus stabilisation remains operational, by evaluating different cylindrical lenses and configurations (Fig. 2a). Calibration curves and time-lapse datasets were acquired to determine operating range and precision for each condition (Fig. 2b). For the 100x 1.35-NA silicone-immersion objective lens, it was possible to increase the operating range from 2.8 μm to 20.6 μm by changing the cylindrical lens from 1000 mm to 300 mm. However, this came at the cost of sensitivity and thus precision that changed from 1.5 nm to 12.7 nm. This means that the performance of the system can be chosen based on the application that is needed by tuning the extent of astigmatism.

**Fig. 2.**
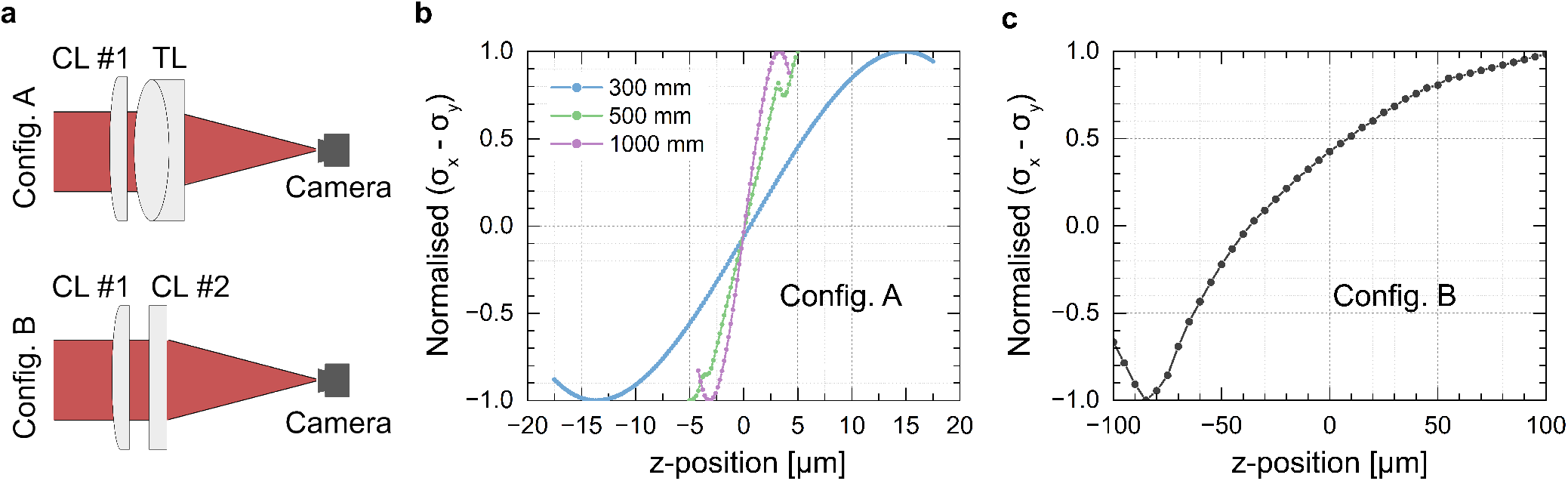
Extending the operating range. (a) Different lens configurations for PiFocus. (b) Calibration using different cylindrical lenses (CL) with 100x 1.35-NA silicone-immersion lens. (c) Calibration curve for two orthogonal cylindrical lenses.

An alternative strategy to further extend the operating range was explored by employing a combination of two cylindrical lenses (Config. B) rather than a CL-TL configuration (Config. A). Results revealed that while this approach greatly increased the operating range >100 μm, it introduced non-linear behaviour in the resulting calibration curve. As the separation between the focal planes is increased, the non-linearity increases, which needs to be taken into account for scanning applications.

### Compatibility with varying NA objective lenses

Conventional TIR-based focus stabilisation, as well as most beam-based methods, are mostly limited to high-NA objective lenses to provide sufficient sensitivity and signal. However, the ability to switch between objective lenses of varying magnifications and NAs while maintaining focus is important to successfully perform multimodal experiments. Furthermore, certain applications require high axial stability but can only be performed with low NA objective lenses. We therefore characterised the performance of PiFocus for a wide variety of commonly used objective lenses. We performed axial drift estimation experiments for each objective lens and demonstrated >10 μm operating range with <21 nm precision for all lenses (Fig. 3a). The results are summarised in the Table shown in Fig. 3b. These results demonstrate that PiFocus offers a universal solution to focus stabilisation, which is critical for microscopes that employ multiple objective lenses with varying NA. The table also includes different cylindrical lens combinations, demonstrating the tunability of astigmatism-based focus stabilisation.

**Fig. 3.**
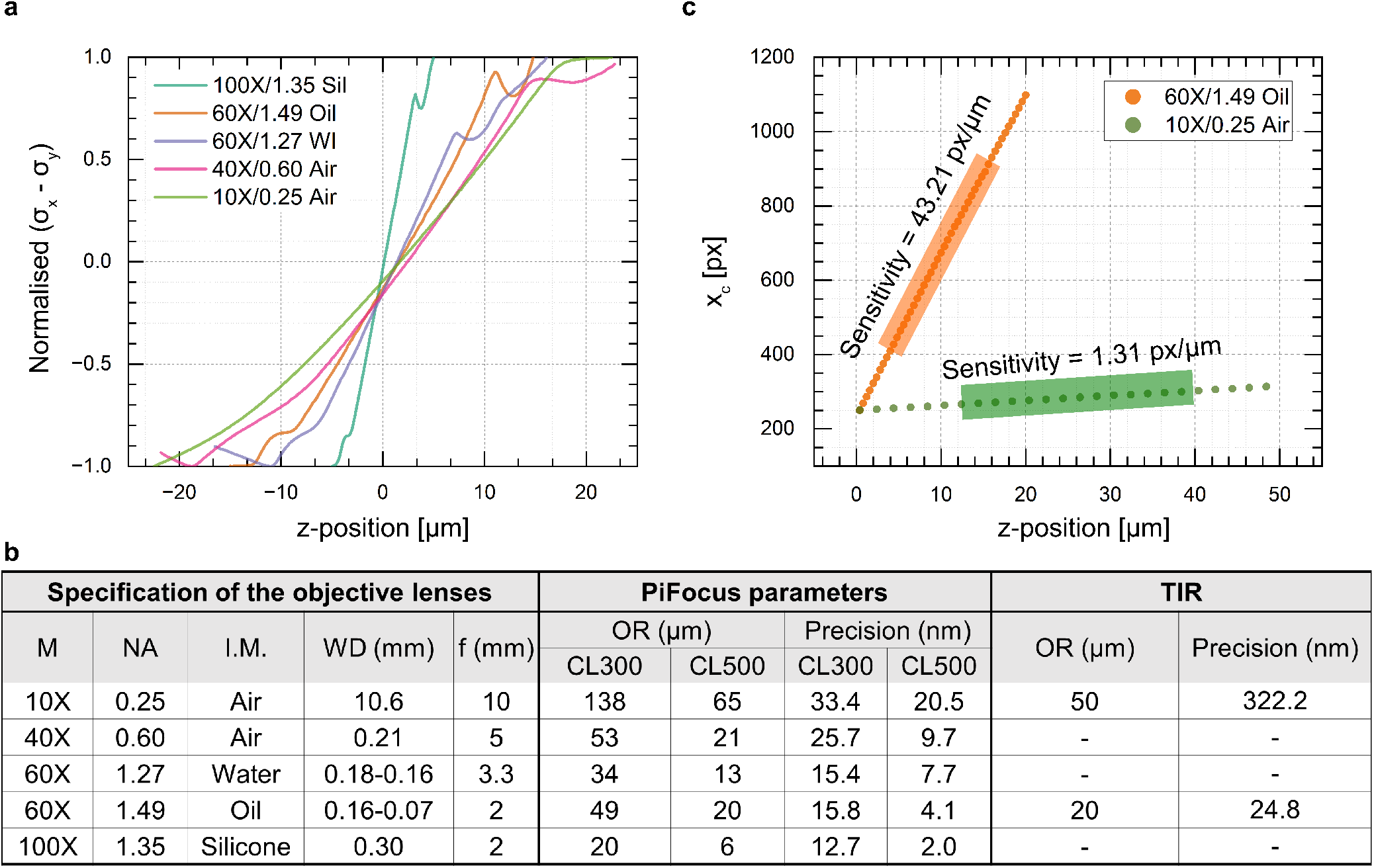
PiFocus offers universal lens compatibility and is superior to TIR for low NA. (a) Calibrations for different objective lenses with varying magnification, NA and immersion media. (b) calibration curve for TIR-based stabilisation performed with 60x 1.49-NA oil-immersion (orange) and 10x 0.25-NA (green) objective lenses. (c) Summary table of all results including comparison between PiFocus and TIR. (M) Magnification, (I.M) Immersion medium, (WD) Working distance, (OR) operating range.

### Comparison with TIR-based focus stabilisation

To demonstrate the universality of PiFocus, we compared its performance to the commonly used TIR-based approach. We implemented TIR stabilisation on the same microscope and determined operating range and precision. The calibrations for the 60x 1.49 NA oil-immersion and 10x 0.25 NA objective lenses are shown in Fig. 3b. The limitations of the TIR-based approach become evident when considering that the sensitivity drops approximately 30-fold upon changing to the low-magnification objective lens, resulting in a precision of 322 nm and an operating range of 50 microns (Fig. 3c, Video S3). Conversely, PiFocus achieves a superior 20.5 nm precision with an operating range of 65 microns. This can be explained by considering that estimation of the main parameters *σ*_*x*_ and *σ*_*y*_ are insensitive to laser pointing stability. As the beam travels ∼2 m to reach the detector, imperfect pointing stability leads to large long-term variations in Xc due to the instability and vibration of optics. This also explains why the performance of our TIR implementation is worse than astigmatism (Fig. 3b) and other similar TIR approaches, which demonstrates the simplicity of implementing astigmatism-based focus stabilisation far away from the sample.

### PiFocus stabilisation

We implemented PiFocus on a Raspberry Pi 4B architecture (Fig. 4a) and custom-written Python scripts were used for image acquisition, calibration and focus stabilisation. The Raspberry Pi interfaced with a piezo z-stage that enabled feedback control of z-position. While in operation, PiFocus continuously acquires images and performs real-time axial drift estimation. Proportional feedback control is then applied to correct the focal position using the piezo stage. To achieve real-time processing at reasonable sampling rates, it was necessary to project images in the x and y dimensions respectively and use 1D Gaussian fitting. ^41^ With this approach, PiFocus was able to maintain the astigmatic beam shape at >100 Hz. By utilising a bypass voltage signal (Fig. 4a) from MicroManager to the Raspberry Pi through an Arduino Uno, PiFocus can perform z-scanning during focus stabilisation.

**Fig. 4.**
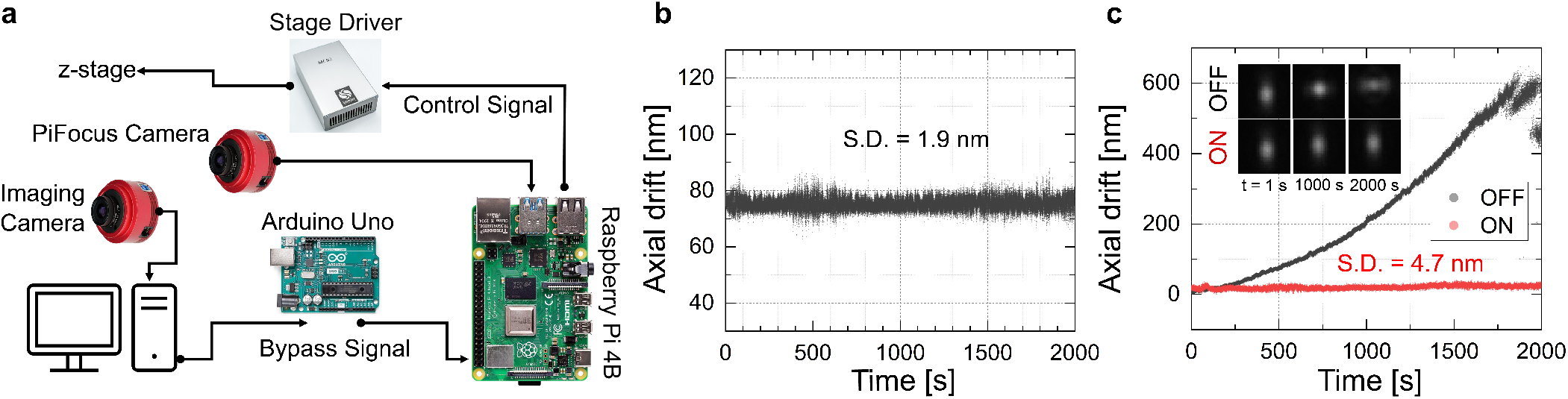
Active focus stabilisation using PiFocus. (a) Schematic of PiFocus hardware implementation. (b) Axial sample drift monitored by PiFocus. (c) Axial drift with stabilisation ON (red) and OFF (black) monitored by imaging fluorescent beads using astigmatic 3D localisation.

Finally, we demonstrated focus stabilisation in a realistic imaging experiment. PiFocus was activated while imaging fluorescent nanoparticles using a fluorescence imaging path that employed astigmatic 3D localisation microscopy (4.4 nm precision, Fig. S3). The astigmatic reference beam with a precision of 1.9 nm was also captured (Fig. 4b). We compared the axial drift of the fluorescent particles with PiFocus on and off (Fig. 4c). Without PiFocus, significant drift (>600 nm over 40 min) could be observed. Conversely with PiFocus activated, the drift was kept below 5 nm, which is at the limit of the axial precision of the fluorescence measurements. These results confirm that the drift estimation experiments done represent the axial response of the imaging system.

## Discussion

We have devised and characterised an active focus stabilisation system, based on real-time monitoring of an astigmatic laser beam reflected from the coverslip-sample interface. Depending on the selected operating range and objective lens, the system achieves sub-10 nm precision in axial drift estimation and correction. This performance is similar to previous TIR-based solutions, although there are a few important differences: (1) We observe good performance (∼20 nm precision) even for low-NA objective lenses, as astigmatism does not require a large NA to achieve high sensitivity. (2) Our approach is not affected by long-term pointing stability variations present in TIR methods, as we use the width rather than the position of the beam. This means that there is no need to keep the focus stabilisation module close to the objective lens, as is usually the case for TIR-methods ^33^ and we demonstrate excellent stability even over extended distances (∼2 m from the fibre output to the beam-monitoring camera). (3) While the operating regime in TIR can be tuned, our approach achieves a significantly higher precision for a given range. (4) Our implementation is carried out on a PI architecture that would also work for a TIR approach and can be independently added on to most existing microscopes.

There are however limitations to our approach. The precision achieved is limited by vibrations, background, and camera noise and astigmatism inherently requires using a camera. It is also necessary to fit a Gaussian function to the data, which is inherently slower than certain centroid methods used to find the beam position. However, we achieve >100 Hz update rates on a relatively slow Raspberry Pi platform, which clearly offers scope for faster CPUs or GPU-acceleration if higher update rates are required. We explored alternative analysis methods based on image moments that were slightly faster, however we found that these were much more susceptive to background fluctuations and thus discarded them.

We demonstrated the tunability of the operating range for focus stabilisation. Through the utilisation of cylindrical lenses featuring different focal lengths, we were able to investigate the fundamental trade-off between the calibration sensitivity and the operating range. As the operating range increases and the distance between the two focal points point is moved, the differing effective NAs give rise to nonlinear behaviour that ultimately affects precision. For deployment on a super-resolution microscope requiring nanometre precision, opting for mild astigmatism with an operating range of ∼10 μm is ideal. Conversely, for imaging larger organisms such as zebrafish, larger separation between the focal points can be employed to extend the operating range beyond 100 μm, albeit with reduced precision exceeding 40 nm and a potentially non-linear calibration. It should be noted that the operating range and/or precision can be further improved by using tunable or translating lenses.

There are similarities between our approach and the recently published openFrame solution. ^32^ The main difference is that openFrame applies astigmatism near the laser launch position that is very mild, which is suitable for large operating ranges. Ultimately, openFrame achieves a precision of 35 nm and a refresh rate of 4 Hz, whereas our approach achieves a precision of down to 2 nm and a refresh rate of 100 Hz, which is more suitable for high-resolution applications.

Our devised solution was implemented on a Raspberry Pi system, providing the capability of serving as an independent focus stabilisation unit on most microscopes that can be turned on with a physical button or via software. The entire system can be implemented for approximately £1,800 (Table S1-2), not including a piezo stage. If an objective piezo scanner is required, it can be sourced for about £2,000. The use of Raspberry Pi is also advantageous as it addresses the difficulty associated with designing and fabricating a bespoke circuit board that is currently required for solutions such as pgFocus. ^30^ While we chose a Raspberry Pi platform here, the astigmatism-based focus stabilisation used in PiFocus could also be integrated with Micromanager, similar to the solution provided for openFrame.

## Author Contributions

A.P. conceptualised, supervised and administered the project. A.R., T.C., A.T.A.A. and A.P. developed the calibration methodology, conducted experiments and analysed data. A.R. designed and implemented the optical setup. A.R. and A.P. carried out the nanoparticle fluorescence measurements. A.R., T.C. and A.P. developed Python code for analysis and focus stabilisation. A.R. and A.P. wrote the manuscript. All authors reviewed and edited the manuscript.

## Acknowledgements

The authors would like to thank summer student Robert Elliott who setup the preliminary camera implementation on Raspberry Pi. We would also like to thank Karl Bellvé for helpful discussion on hardware. This work was funded by a University of Leeds University Academic Fellowship, a Royal Society Research Grant (RGS*\*R2*\*202446), as well as an AMS Springboard Award (SBF006*\*1138), awarded to A.P., and by an EPSRC Bragg Centre Doctoral Training Studentship (EP/T517860/1) awarded to A.R.

## Methods

### Optical setup

A diagram of the optical setup is shown in Fig. S4. The focus stabilisation system used a 300 mW, 808 nm multimode laser diode (OFL311, OdicForce Lasers) with 9 nm bandwidth. The output beam of this laser was collimated with a triplet lens assembly (OFL177, OdicForce) and the beam was then coupled into a 780-970 nm single-mode fibre (P1-780A-FC-1, Thorlabs) using a 10x 0.25 NA objective lens (Olympus), providing an output power of 2.6 mW. Some experiments (TIR, 40x and 10x) also used a 700 mW, 638 nm multimode laser diode (OFL195-U, OdicForce), although the results were identical for the two lasers (Fig. S2). The output of this fibre was collimated using a short focal length lens (f = 40 mm, AC254-040-B, Thorlabs). An adjustable iris (ID25SS/M, Thorlabs) was utilised to adjust the beam size. The collimated beam with a 5 mm diameter then passed a 50/50 cube beam splitter (CCM1-BS014/M, Thorlabs). A polarising beam splitter could be used instead to increase signal and to minimise reflections. The beam entered the imaging path by being reflected from the NIR shortpass dichroic (#69-196, Edmund Optics) heading towards the back aperture of the microscope objective lens. The laser beam was then focused by the imaging objective lens onto the top of the #1.5 glass coverslip. The back-reflected light from the coverslip-sample interface returned to the beam splitter and was partially reflected onto the camera detector. The reflection was filtered by a NIR longpass filter (FELH0800, Thorlabs), passed through a cylindrical lens (with varying focal lengths), which was positioned after the beam splitter, 40 mm before the tube lens (f = 200, TTL200-B, Thorlabs). The camera was placed at the centre of the two focal points created by the cylindrical lens and the tube lens. Two cameras were explored, a low-cost (∼£60) high-speed 10-bit Raspberry Pi camera (Arducam OV9782) and a medium-cost (∼£300) astronomy camera based on the IMX290 sensor (ZWO ASI290MM). Both cameras performed similarly in terms of speed and precision, although the ZWO camera was more versatile and offered higher bit depth. In an alternative configuration, the tube lens was replaced by a second orthogonal cylindrical lens to attempt to minimise non-linear behaviour in the calibration curves.

The focus stabilisation unit was integrated into a custom-built fluorescence microscope capable of performing epi-, Total Internal Reflection (TIR), ^46^ and Highly Inclined and Laminated Optical sheet (HiLo) ^47^ excitation for imaging. A three-axis nanopositioning piezo stage (SLC-1780-D-S, SmarAct), equipped with an integrated sensors for closed-loop positioning, was used to control the sample position. ^33^

### Fluorescence setup

For fluorescence excitation, a 4.5 mW 639 nm collimated laser module (PL252, Thorlabs) was used, which was passed through an excitation filter and focused onto the back focal plane of the objective lens using an achromatic lens (AC254-200-A-ML, Thorlabs). With a mirror and the lens on a translational stage, we were able to switch from widefield to HiLo and TIR excitation by translating the laser beam toward the periphery of the back focal plane of the objective lens. The excitation laser beam was then delivered to the objective lens using a quadband dichroic mirror (Di03-R405/488/561/635-t3-25x36, Semrock). The emitted light by the fluorescent sample was collected by the objective lens and passed through the dichroic mirror, an emission (FELH0650, Thorlabs) and a bandpass filter (FF01-609/181-25, Semrock) was used to block the NIR beam. A tube lens (TTL200-A, f = 200 mm, Thorlabs) was finally used to create an image of the focal plane on a CMOS camera (ASI485MC, ZWO).

### Calibration and precision measurements

A z-image stack was acquired to establish a calibration curve that maps the relationship between the focal plane position and the corresponding beam shape, which was used for focus stabilisation. The z-stack scan range and step size were chosen to ensure that the entire range of astigmatism was covered, encompassing the full range of potential focal plane deviations. For each frame, the average intensity in x and y (2D image projected into a 1D curve) were calculated and fitted with a Gaussian function to determine *σ*_*x*_ and *σ*_*y*_.

To determine the precision of the z-position estimation, a time-lapse image stack was acquired with 200 fps frame rate for 100 seconds. The precision was then calculated as the standard deviation of *σ*_*y*_ − *σ*_*x*_. Data acquisition and instrument control were performed using a custom-written Python script. Data analysis and processing were performed with scripts written in Python. All scripts are provided on Github. ^48^

### Fiducial benchmark

To evaluate the performance of the focus stabilisation unit to maintain focus, we monitored the position of a fluorescent nanoparticle in 3D over time. We used 200 nm FluoSpheres (F8807, Thermo Fisher) as the fiducial marker. ^49^ To acquire astigmatic single-molecule images of the fluorescent beads, we introduced weak astigmatism by placing a long focal length cylindrical lens (f = 1000 mm, LJ1516RM-A, Thorlabs) in front of the imaging camera. We acquired a 3 μm z-stack to generate a calibration curve (Fig. S3) for the beads. After activating PiFocus, we started a long time-lapse acquisition (1 hr) of the selected bead with 40 ms exposure time. We then turned the stabilisation system off and captured another long time-lapse acquisition (1 hr). The image acquisition was performed using MicroManager 2.0. The calibration stack and time-lapses were analysed using PeakFit, ^50^ to extract *σ*_*y*_ −*σ*_*x*_ for z-position and precision estimations, as done for the focus stabilisation.

### Raspberry Pi implementation

The focus stabilisation system was implemented on a Raspberry Pi 4 Model B (OKdo 4GB starter kit). The Arducam was connected through the Raspberry Pi camera flex cable and the ZWO and the piezo stage driver were connected to the USB 3.0 port of the Raspberry Pi board. Focus stabilisation was achieved through Python scripts available on Github. ^48^ A polynomial, corresponding to the calibration for the given objective lens, is used to estimate the current and target positions. For each frame, fitting is performed as described previously to determine *σ*_*y*_ − *σ*_*x*_. The offset from the target position is then calculated based on the calibration and proportional control is applied to maintain focus. In our case we had direct USB control over the piezo stage. For more conventional solutions, the piezo could be controlled using a DAC voltage signal, that can be achieved using a 16-bit DAC (AD5693, Adafruit) and we have included code for this. To enable z-scanning during focus stabilisation, we used an Arduino Uno that interfaced directly with Micromanager. The Arduino outputs a bypass voltage through the AD5693 16-bit DAC that is read by a 16-bit ADC (ADS1115, Adafruit) on the Raspberry Pi. When a change in the bypass voltage is detected, the piezo stage is moved to the new target position and the *σ*_*y*_ −*σ*_*x*_ setpoint is modified based on the calibration curve to maintain stabilisation during scanning.

## Bibliography

1. B. Neumann, A. Dämon, D. Hogenkamp, E. Beckmann, and J. Kollmann, “A laser-autofocus for automatic microscopy and metrology,” Sensors and Actuators, vol. 17, no. 1-2, pp. 267–272, 1989.

2. S. A. Jones, S.-H. Shim, J. He, and X. Zhuang, “Fast, three-dimensional super-resolution imaging of live cells,” Nature Methods, vol. 8, no. 6, pp. 499–505, 2011.

3. T. Orré, A. Joly, Z. Karatas, B. Kastberger, C. Cabriel, R. T. Böttcher, S. Lévêque-Fort, J.-B. Sibarita, R. Fässler, B. Wehrle-Haller, O. Rossier, and G. Giannone, “Molecular motion and tridimensional nanoscale localization of kindlin control integrin activation in focal adhesions,” Nature Communications, vol. 12, no. 1, p. 3104, 2021.

4. P. Jouchet, C. Cabriel, N. Bourg, M. Bardou, C. Poüs, E. Fort, and S. Lévêque-Fort, “Nanometric axial localization of single fluorescent molecules with modulated excitation,” Nature Photonics, 2021.

5. M. Bates, J. Keller-Findeisen, A. Przybylski, A. Hüper, T. Stephan, P. Ilgen, A. R. Cereceda Delgado, E. D’Este, A. Egner, S. Jakobs, S. J. Sahl, and S. W. Hell, “Optimal precision and accuracy in 4Pi-STORM using dynamic spline PSF models,” Nature Methods, 2022.

6. E. W. Sanders, A. R. Carr, E. Bruggeman, M. Körbel, S. I. Benaissa, R. F. Donat, A. M. Santos, J. McColl, K. O’Holleran, D. Klenerman, S. J. Davis, S. F. Lee, and A. Ponjavic, “resPAINT: Accelerating volumetric super-resolution localisation microscopy by active control of probe emission,” Angewandte Chemie International Edition, p. anie.202206919, 2022.

7. S. C. M. Reinhardt, L. A. Masullo, I. Baudrexel, P. R. Steen, R. Kowalewski, A. S. Eklund, S. Strauss, E. M. Unterauer, T. Schlichthaerle, M. T. Strauss, C. Klein, and R. Jungmann, “Ångström-resolution fluorescence microscopy,” Nature, vol. 617, no. 7962, pp. 711–716, 2023.

8. M. Klevanski, F. Herrmannsdoerfer, S. Sass, V. Venkataramani, M. Heilemann, and T. Kuner, “Automated highly multiplexed super-resolution imaging of protein nano-architecture in cells and tissues,” Nature Communications, vol. 11, no. 1, p. 1552, 2020.

9. R. Diekmann, M. Kahnwald, A. Schoenit, J. Deschamps, U. Matti, and J. Ries, “Optimizing imaging speed and excitation intensity for single-molecule localization microscopy,” Nature Methods, vol. 17, no. 9, pp. 909–912, 2020.

10. A. E. S. Barentine, Y. Lin, E. M. Courvan, P. Kidd, M. Liu, L. Balduf, T. Phan, F. Rivera-Molina, M. R. Grace, Z. Marin, M. Lessard, J. Rios Chen, S. Wang, K. M. Neugebauer, J. Bewersdorf, and D. Baddeley, “An integrated platform for high-throughput nanoscopy,” Nature Biotechnology, 2023.

11. R. M. Power, A. Tschanz, T. Zimmermann, and J. Ries, “Automated 3D multi-color single-molecule localization microscopy,” preprint, Biophysics, 2023.

12. G. Reinheimer, “Anordnung zum selbsttätigen Fokussieren auf in optischen Geräten zu betrachtende Objekte,” 1971.

13. Q. Li, “Autofocus system for microscope,” Optical Engineering, vol. 41, no. 6, p. 1289, 2002.

14. Nikon, “Perfect Focus System.” https://www.nikonusa.com/fileuploads/pdfs/Instruments/TE2000_PFS.pdf.

15. Zeiss, “Definite Focus.” https://www.imdik.pan.pl/images/Us%C5%82ugi_dokumenty/Przydatne_informacje/2d_Definite_Focus_information.pdf.

16. L. Microsystems, “Leica Adaptive Focus Control.” https://www.leica-microsystems.com/products/light-microscopes/p/leica-dmi6000-with-adaptive-focus-control/.

17. Olympus, “TruFocus: Z Drift Compensator.” https://www.olympus-lifescience.com/en/microscopes/inverted/ix83/trufocus/.

18. W. Hwang, S. Bae, and S. Hohng, “Autofocusing system based on optical astigmatism analysis of single-molecule images,” Optics Express, vol. 20, no. 28, p. 29353, 2012.

19. P. Bon, N. Bourg, S. Lécart, S. Monneret, E. Fort, J. Wenger, and S. Lévêque-Fort, “Three-dimensional nanometre localization of nanoparticles to enhance super-resolution microscopy,” Nature Communications, vol. 6, no. 1, p. 7764, 2015.

20. R. Schmidt, T. Weihs, C. A. Wurm, I. Jansen, J. Rehman, S. J. Sahl, and S. W. Hell, “MINFLUX nanometer-scale 3D imaging and microsecond-range tracking on a common fluorescence microscope,” Nature Communications, vol. 12, no. 1, p. 1478, 2021.

21. Y. Liron, Y. Paran, N. G. Zatorsky, B. Geiger, and Z. Kam, “Laser autofocusing system for high-resolution cell biological imaging,” Journal of Microscopy, vol. 221, no. 2, pp. 145–151, 2006.

22. P. Kner, J. Sedat, D. Agard, and Z. Kam, “High-resolution wide-field microscopy with adaptive optics for spherical aberration correction and motionless focusing,” Journal of Microscopy, vol. 237, no. 2, pp. 136–147, 2010.

23. A. Pertsinidis, Y. Zhang, and S. Chu, “Subnanometre single-molecule localization, registration and distance measurements,” Nature, vol. 466, no. 7306, pp. 647–651, 2010.

24. C.-S. Liu, Y.-C. Lin, and P.-H. Hu, “Design and characterization of precise laser-based autofocusing microscope with reduced geometrical fluctuations,” Microsystem Technologies, vol. 19, no. 11, pp. 1717–1724, 2013.

25. B. X. Cao, P. Hoang, S. Ahn, J.-o. Kim, H. Sohn, and J. Noh, “Realtime detection of focal position of workpiece surface during laser processing using diffractive beam samplers,” Optics and Lasers in Engineering, vol. 86, pp. 92–97, 2016.

26. C. Niederauer, M. Seynen, J. Zomerdijk, M. Kamp, and K. A. Ganzinger, “The K2: Open-source simultaneous triple-color TIRF microscope for live-cell and single-molecule imaging,” HardwareX, vol. 13, p. e00404, 2023.

27. B. Hajj, J. Wisniewski, M. El Beheiry, J. Chen, A. Revyakin, C. Wu, and M. Dahan, “Whole-cell, multicolor superresolution imaging using volumetric multifocus microscopy,” Proceedings of the National Academy of Sciences, vol. 111, no. 49, pp. 17480–17485, 2014.

28. N. Saliba, G. Gagliano, and A.-K. Gustavsson, “Whole-cell multi-target single-molecule super-resolution imaging in 3D with microfluidics and a single-objective tilted light sheet,” preprint, Biophysics, 2023.

29. S. Coelho, J. Baek, J. Walsh, J. J. Gooding, and K. Gaus, “3D active stabilization for single-molecule imaging,” Nature Protocols, vol. 16, no. 1, pp. 497–515, 2021.

30. K. Bellve, C. Standley, L. Lifshitz, and K. Fogarty, “Design and Implementation of 3D Focus Stabilization for Fluorescence Microscopy,” Biophysical Journal, vol. 106, no. 2, p. 606a, 2014.

31. K. Bellve, C. Standley, L. Lifshitz, and K. Fogarty, “pgFocus: wiki page.” https://big.umassmed.edu/wiki/index.php/PgFocus.

32. J. Lightley, S. Kumar, M. Q. Lim, E. Garcia, F. Görlitz, Y. Alexandrov, T. Parrado, C. Hollick, E. Steele, K. Roßmann, J. Graham, J. Broichhagen, I. A. McNeish, C. A. Roufosse, M. A. A. Neil, C. Dunsby, and P. M. W. French, “openFrame : A modular, sustainable, open microscopy platform with single-shot, dual-axis optical autofocus module providing high precision and long range of operation,” Journal of Microscopy, p. jmi.13219, 2023.

33. J. S. H. Danial, J. Y. L. Lam, Y. Wu, M. Woolley, E. Dimou, M. R. Cheetham, D. Emin, and D. Klenerman, “Constructing a cost-efficient, high-throughput and high-quality single-molecule localization microscope for super-resolution imaging,” Nature Protocols, vol. 17, no. 11, pp. 2570–2619, 2022.

34. B. Van Den Berg, R. Van Den Eynde, B. Amouroux, M. Müller, P. Dedecker, and W. Vandenberg, “A Modular Approach to Active Focus Stabilization for Fluorescence Microscopy,” preprint, Biochemistry, 2020.

35. B. Cao, P. Hoang, S. Ahn, J.-o. Kim, H. Kang, and J. Noh, “In-Situ Real-Time Focus Detection during Laser Processing Using Double-Hole Masks and Advanced Image Sensor Software,” Sensors, vol. 17, no. 7, p. 1540, 2017.

36. M. Bathe-Peters, P. Annibale, and M. J. Lohse, “All-optical microscope autofocus based on an electrically tunable lens and a totally internally reflected IR laser,” Optics Express, vol. 26, no. 3, p. 2359, 2018.

37. S.-Y. Chen, R. Heintzmann, and C. Cremer, “Sample drift estimation method based on speckle patterns formed by backscattered laser light,” Biomedical Optics Express, vol. 10, no. 12, p. 6462, 2019.

38. L. Silvestri, M. C. Müllenbroich, I. Costantini, A. P. Di Giovanna, G. Mazzamuto, A. Franceschini, D. Kutra, A. Kreshuk, C. Checcucci, L. O. Toresano, P. Frasconi, L. Sacconi, and F. S. Pavone, “Universal autofocus for quantitative volumetric microscopy of whole mouse brains,” Nature Methods, vol. 18, no. 8, pp. 953–958, 2021.

39. H. Ma and Y. Liu, “Embedded nanometer position tracking based on enhanced phasor analysis,” Optics Letters, vol. 46, no. 16, p. 3825, 2021.

40. D. K. Cohen, W. H. Gee, M. Ludeke, and J. Lewkowicz, “Automatic focus control: the astigmatic lens approach,” Applied Optics, vol. 23, no. 4, p. 565, 1984.

41. W.-Y. Hsu, C.-S. Lee, P.-J. Chen, N.-T. Chen, F.-Z. Chen, Z.-R. Yu, C.-H. Kuo, and C.-H. Hwang, “Development of the fast astigmatic auto-focus microscope system,” Measurement Science and Technology, vol. 20, no. 4, p. 045902, 2009.

42. J. Lightley, F. Görlitz, S. Kumar, R. Kalita, A. Kolbeinsson, E. Garcia, Y. Alexandrov, V. Bousgouni, R. Wysoczanski, P. Barnes, L. Donnelly, C. Bakal, C. Dunsby, M. A. Neil, S. Flaxman, and P. M. French, “Robust deep learning optical autofocus system applied to automated multiwell plate single molecule localization microscopy,” Journal of Microscopy, vol. 288, no. 2, pp. 130–141, 2022.

43. J. Ortega Arroyo, D. Cole, and P. Kukura, “Interferometric scattering microscopy and its combination with single-molecule fluorescence imaging,” Nature Protocols, vol. 11, no. 4, pp. 617–633, 2016.

44. S. Lin, Y. He, D. Feng, M. Piliarik, and X.-W. Chen, “Optical Fingerprint of Flat Substrate Surface and Marker-Free Lateral Displacement Detection with Angstrom-Level Precision,” Physical Review Letters, vol. 129, no. 21, p. 213201, 2022.

45. R. Chowdhury, A. Sau, J. Chao, A. Sharma, and S. M. Musser, “Tuning axial and lateral localization precision in 3D super-resolution microscopy with variable astigmatism,” Optics Letters, vol. 47, no. 21, p. 5727, 2022.

46. D. Axelrod, “Cell-substrate contacts illuminated by total internal reflection fluorescence.,” Journal of Cell Biology, vol. 89, no. 1, pp. 141–145, 1981.

47. M. Tokunaga, N. Imamoto, and K. Sakata-Sogawa, “Highly inclined thin illumination enables clear single-molecule imaging in cells,” Nature Methods, vol. 5, no. 2, pp. 159–161, 2008.

48. A. Rahmani, T. Cox, and A. Ponjavic, “PiFocus: Acquisition, Analysis and Hardware Control,” 2023. https://github.com/PonjavicLab/PiFocus.

49. A. Balinovic, D. Albrecht, and U. Endesfelder, “Spectrally red-shifted fluorescent fiducial markers for optimal drift correction in localization microscopy,” Journal of Physics D: Applied Physics, vol. 52, no. 20, p. 204002, 2019.

50. T. J. Etheridge, A. M. Carr, and A. D. Herbert, “GDSC SMLM: Single-molecule localisation microscopy software for ImageJ,” Wellcome Open Research, vol. 7, p. 241, 2022.

