## Supplementary Document for "Astigmatism-based focus stabilisation with universal objective lens compatibility, extended operating range and nanometre precision"

### Supplementary Figures

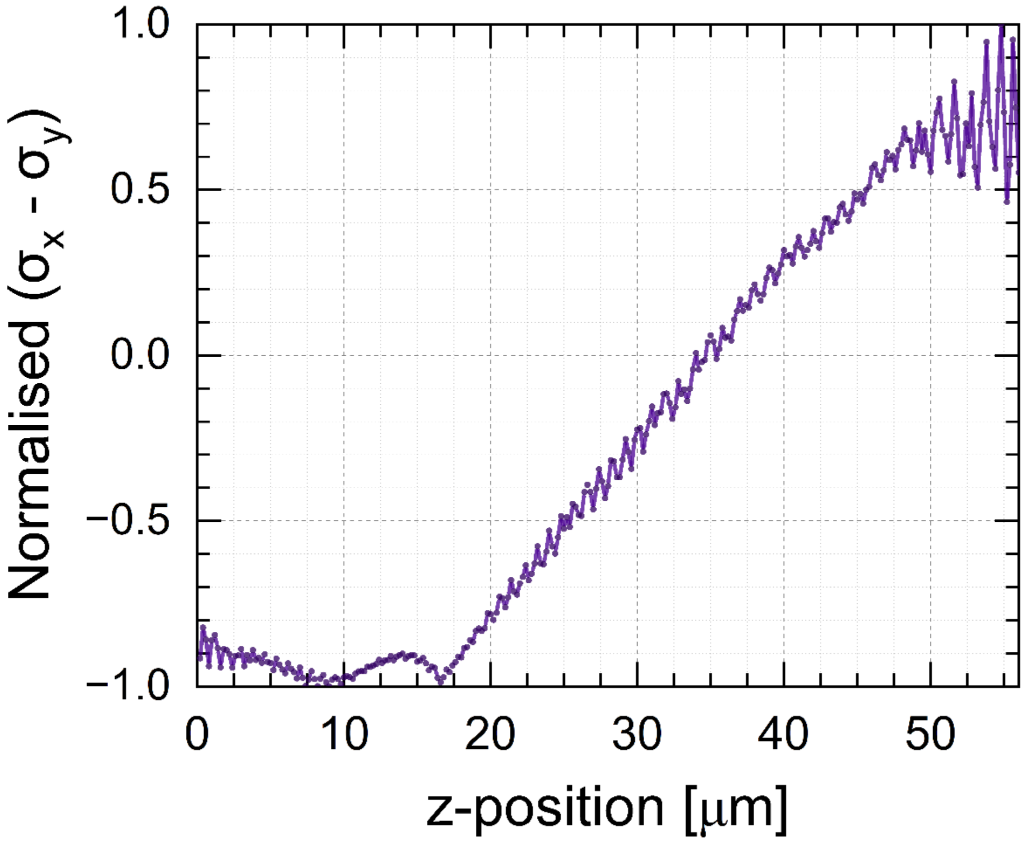


**Figure S1:** Calibration curve of the PiFocus system with 40X/0.40 objective lens before removing the interference pattern.

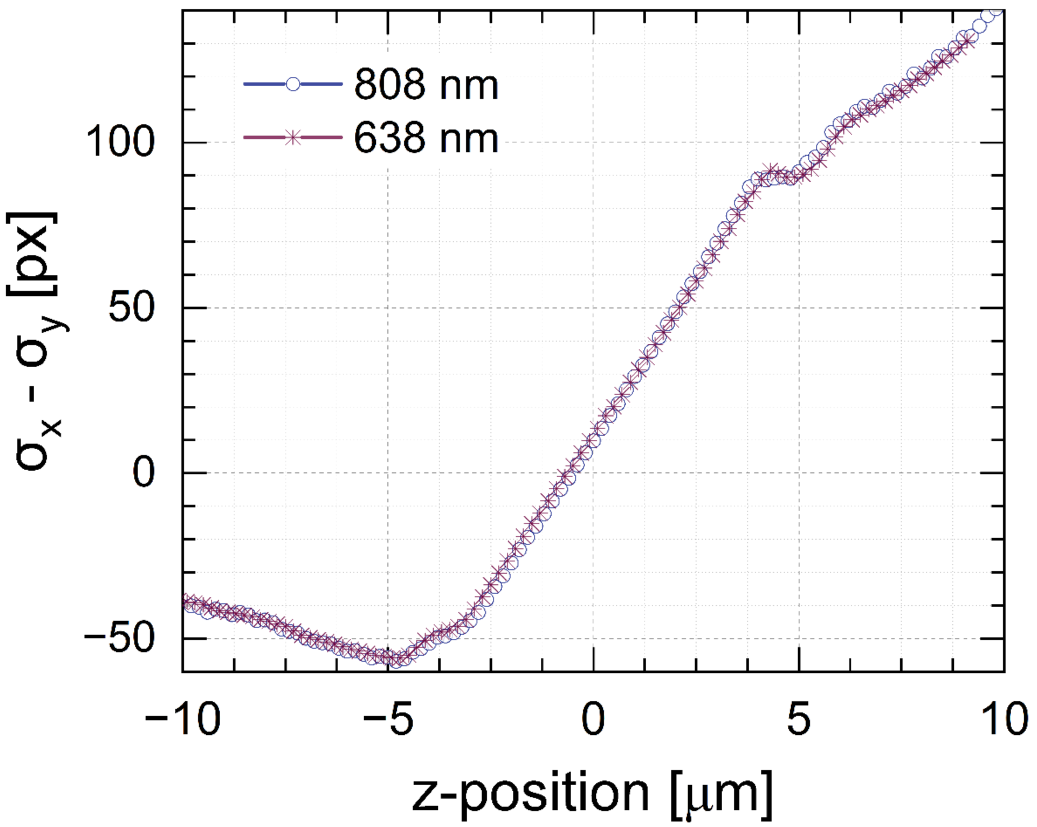


**Figure S2:** Overlap of the calibration curves of the PiFocus system for 808 nm and 638 nm lasers used for experiments.

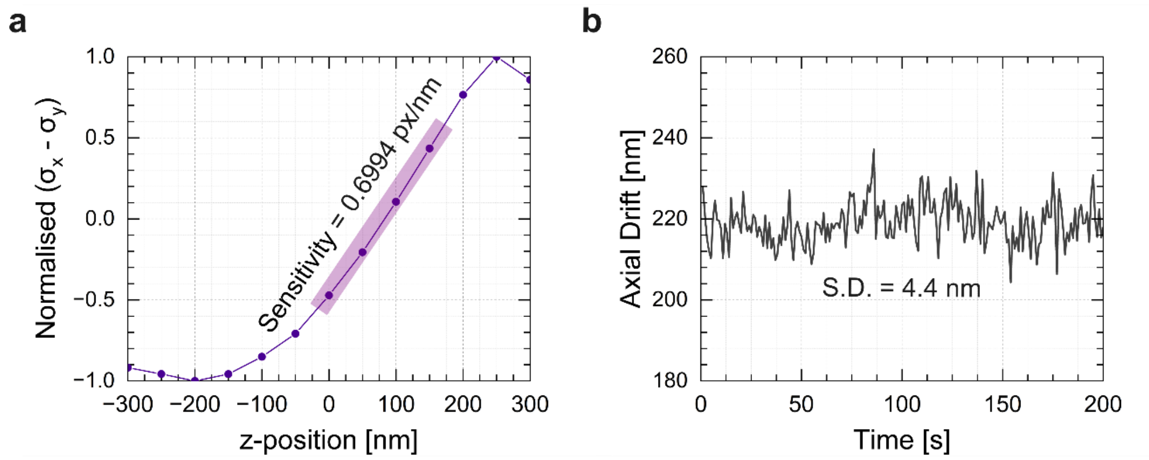


**Figure S3:** ((a) Calibration curve for the fluorescence 3D astigmatism setup. (b) Timelapse for a fiducial marker at z = 0.

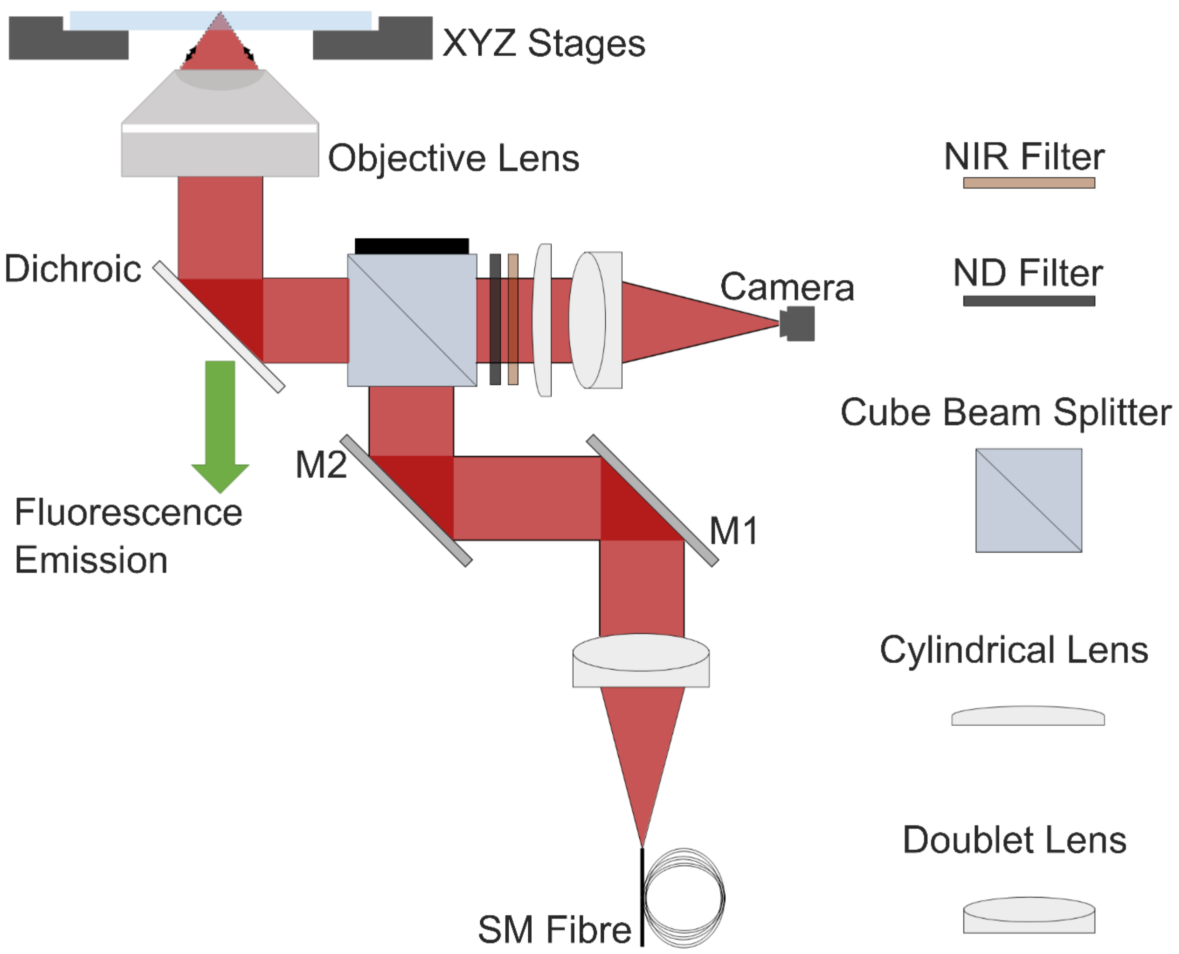


**Figure S4:** Schematic of PiFocus optical path.

**Table S1:** Optics and optomechanics used for PiFocus

| Item | Part code | Company | Price | Quantity |
|---|---|---|---|---|
| Laser Diode Housing | OFL448 | OdicForce Lasers | £9 | 1 |
| Laser Diode | OFL311 | OdicForce Lasers | £5 | 1 |
| Collimating lens | OFL177 | OdicForce Lasers | £5 | 1 |
| Focus Ring Holder | OFL458 | OdicForce Lasers | £2 | 1 |
| Fibre coupling lens | PA10X-INF | AmScope | £40 | 1 |
| Fibre Adapter | SM1FCA | Thorlabs | £30 | 2 |
| Single-mode fiber | P1-780A-FC-1 | Thorlabs | £80 | 1 |
| Collimating Lens | AC254-040-B-ML | Thorlabs | £106 | 1 |
| Adjustable Iris | ID25SS/M | Thorlabs | £56 | 1 |
| Mirrors | BB1-E03 | Thorlabs | £65 | 2 |
| Cube Beam Splitter | CCM1-BS014/M | Thorlabs | £270 | 1 |
| Polarising Beam Splitter | CCM1-PBS255/M | Thorlabs | £290 | 1 |
| IR Shortpass Dichroic | #69-196 | Edmund Optics | £107 | 1 |
| IR Longpass Filter | FELH0800 | Thorlabs | £115 | 1 |
| Cylindrical Lens | LJ1558RM-B | Thorlabs | £105 | 1 |
| Cylindrical Lens | LJ1144RM-B | Thorlabs | £113 | 1 |
| Cylindrical Lens | LJ1516RM-B | Thorlabs | £113 | 1 |
| Tube Lens | TTL200-B | Thorlabs | £428 | 1 |
| Alternative tube lens | LA1979-B-ML | Thorlabs | £58 | 1 |
| 3-axis Piezo Stage | SLC-1780-D-S | SmarAct | €9000 | 1 |
| Alternative Piezo Stage | P72.Z100 and E53.B1S | CoreMorrow | $2200 | 1 |

**Table S2:** Electronics and cameras used for PiFocus

| Item | Part code | Company | Price | Quantity |
|---|---|---|---|---|
| Camera | ASI290MM | ZWO ASI | £239 | 1 |
| Camera | OV9782 | Arducam | £55 | 1 |
| Raspberry Pi 4 | Model 4B 4GB | OKdo | £100 | 1 |
| DAC | AD5693R | Adafruit | £9 | 1 |
| ADC | ADS1115 | Adafruit | £11 | 1 |
| Arduino Uno | Uno Rev 3 | Arduino | £20 | 1 |

### Supplementary Videos

**Video S1:** Calibration and time-lapse for the 100x 1.35 silicone-immersion objective lens (10 fps playback speed, Fire LUT).

**Video S2:** A calibration scan that shows the interference pattern created from a coherent laser diode (10 fps playback speed, 60x 1.27 water-Immersion objective lens, Fire LUT).

**Video S3:** TIR Scan for 60x 1.49 oil-immersion and 10x 0.25 Air objective lenses with 500 nm steps (10 fps playback speed, Fire LUT).
